# Nonmuscle myosin 2 turnover in cells is synergistically controlled by the tail and the motor domain

**DOI:** 10.1101/2025.04.30.651508

**Authors:** Anil Chougule, Tatyana M. Svitkina

## Abstract

Myosin 2, an actin-dependent motor, is universally responsible for cell contractility due to its ability to form bipolar filaments. Fast turnover of nonmuscle myosin 2 (NM2) filaments is necessary to keep up with cell motility and shape changes. The turnover mechanisms are not fully understood and differ for two main mammalian paralogs—NM2A and NM2B—whereas paralog copolymerization adds complexity to this process by enabling the intrinsically fast NM2A to dynamize the intrinsically slow NM2B. Here, we show that the nonhelical tail, the C-terminal phosphorylation sites, and surprisingly, the motor domain of the NM2A heavy chain synergistically accelerate the turnover of NM2B in trans and cell motility, suggesting that these three mechanisms collectively control NM2A’s own dynamics. Conversely, the phosphomimetic NM2B tail facilitates only local turnover of endogenous wild type NM2B but not its global redistribution unless the NM2A motor is combined with the phosphomimetic NM2B tail. Collectively, we reveal the cooperation between the motor activity and the NM2 tail-targeting turnover mechanisms in regulating the NM2 filament turnover in trans and cell motility.

**SUMMARY:** Turnover of nonmuscle myosin 2 (NM2) filaments is essential for cell motility and is regulated by various mechanisms including copolymerization of the NM2A and NM2B paralogs. Chougule and Svitkina reveal how the intrinsically fast NM2A accelerates dynamics of slower NM2B in trans.

## INTRODUCTION

Actin-dependent motors of myosin 2 family are uniquely responsible for generation of contractile force in cells due to polymerization into bipolar filaments. Members of nonmuscle myosin 2 (NM2) subfamily are ubiquitous and play essential but diverse roles in all cells, in both healthy and diseased conditions (Chinthalapudi and Heissler, 2024; Garrido-Casado et al., 2024; Quintanilla et al., 2023; Shutova and Svitkina, 2018). In contrast to the striated muscle paralogs, NM2 makes highly dynamic filaments capable of keeping up with cell motility and shape changes. Many NM2 mutations, including those affecting bipolar filament turnover, are associated with human disorders (Asensio-Juarez et al., 2020).

Individual NM2 molecules are hexamers consisting of two heavy chains, each comprising an N-terminal motor domain, a long α-helical rod responsible for heavy chain dimerization via coiled coil formation, and C-terminal nonhelical tailpiece (NHT), as well as two pairs of light chains. Upon phosphorylation of the regulatory light chain, autoinhibited NM2 molecules unfold and acquire both the motor and polymerization activities. In migrating cells, newly assembled NM2 filaments often form clusters and stacks associated with misaligned actin filaments (Quintanilla et al., 2024; Svitkina et al., 1997; Verkhovsky et al., 1995). These clusters then can reorganize into stress fibers – aligned bundles of actin and NM2 filaments – in a process largely driven by NM2 (Beach et al., 2017; Fenix et al., 2016; Hu et al., 2017; Lehtimaki et al., 2021). Whereas the assembly mechanisms of stress fibers in nonmuscle cells are relatively well studied, much less is known how NM2 bipolar filaments undergo paralog-specific depolymerization and recycling to allow reorganization of the contractile system.

Mammalian cells express three different NM2 heavy chains (NM2A, B and C, encoded by *MYH9, MYH10* and *MYH14*, respectively), which have both unique and overlapping roles in cells (Chinthalapudi and Heissler, 2024; Sellers and Heissler, 2019; Shutova and Svitkina, 2018). NM2A and NM2B are widely expressed, whereas NM2C expression is more limited. NM2A is a faster motor than NM2B, whereas NM2B is more processive with stronger catch-bond behavior (Melli et al., 2018; Sellers and Heissler, 2019). In cells, NM2A undergoes faster turnover than NM2B (Raab et al., 2012; Sandquist and Means, 2008; Vicente-Manzanares et al., 2008; Weissenbruch et al., 2022) and is more evenly distributed in migrating cells, whereas NM2B is shifted away from the leading edge (Kolega, 2003; Maupin et al., 1994; Shutova et al., 2017). During cell migration, NM2B is thought to prevent protrusion at the rear, whereas NM2A can quickly redistribute to new protrusions to support traction for the forward advance.

Despite these differences, NM2A and NM2B are sufficiently similar to copolymerize in cells (Beach et al., 2014; Shutova et al., 2014). Moreover, copolymerization of NM2A and NM2B, together with their differential dynamics, explains the front-rear polarization of NM2A and NM2B through self-organization (Shutova et al., 2017; Shutova and Svitkina, 2018). In this mechanism, the cycles of preferential dissociation of NM2A subunits from heterotypic NM2 filaments followed by non-selective recruitment of new subunits gradually produces NM2B-enriched filaments that are more stable and become shifted toward the cell rear by retrograde flow. At the higher NM2A:NM2B ratio, NM2A accelerates the otherwise slow dynamics of NM2B filaments (Shutova et al., 2017) and decreases the cytoskeleton-associated fraction of NM2B (Schiffhauer et al., 2019), because when NM2A subunits dissociate from heterofilaments, the few remaining NM2B subunits cannot stay in a filamentous form and are forced to recycle as well (Shutova and Svitkina, 2018).

Given the key roles of the different disassembly rates of NM2A and NM2B for the organization of the cellular contractile system, it is important to know the underlying mechanisms. Available data indicate that NM2 depolymerization mechanisms primarily target the C-terminal region of the heavy chain, which includes the NHT, contains phosphorylation sites, and binds regulatory proteins (Chinthalapudi and Heissler, 2024; Dulyaninova and Bresnick, 2013; Shutova and Svitkina, 2018). Each of these mechanisms can individually enhance intracellular NM2A dynamics (Breckenridge et al., 2009; West-Foyle et al., 2018), but it remains unclear whether they act redundantly or cooperatively. For NM2B, the C-terminal phosphorylation is a primary mechanism of accelerating its dynamics (Juanes-Garcia et al., 2015). Whether different motor activities of NM2A and NM2B regulate their turnover is not clear.

In this study, we systematically analyzed what features of NM2 heavy chains contribute to their ability to dynamize and redistribute the wild type NM2B in trans. The rationale is that ectopic NM2A with impaired depolymerization would be unable to redistribute endogenous NM2B through copolymerization. Conversely, exogenous NM2B with an improved turnover due to a phosphomimetic mutation would redistribute endogenous wild type NM2B akin to NM2A. As a test system, we use COS7 cells, which lack endogenous NM2A, so that expression of exogenous NM2A readily redistributes endogenous NM2B (Shutova et al., 2017). Using various NM2A mutants, we show that the NHT, the C-terminal phosphorylation sites and, surprisingly, the motor domain synergistically enable NM2A to dynamize NM2B, whereas the dynamizing S1935D mutation in the tail of exogenous NM2B only partially mimics the NM2A effect unless it is combined with the NM2A motor. Collectively, we reveal the cooperation between the motor activity and the two NM2 tail-targeting turnover mechanisms in regulation of the NM2 filament turnover in trans.

## RESULTS

In COS7 cells, which lack endogenous NM2A, the actin-NM2B contractile system consists of a few sharply defined, largely disconnected and barely dynamic ventral stress fibers, while expression of NM2A converts them into a more polymorphic and dynamic system, where endogenous NM2B acquires a more homogenous distribution with enhanced turnover (Shutova et al., 2017). The fast dynamics of NM2A is at the core of the mechanism by which NM2A accelerates turnover of NM2B. Building on this phenomenon, we designed a strategy to reveal the NM2A determinants that control its own fast turnover and, therefore, the reorganization of NM2B in trans. Namely, we expressed in COS7 cells the NM2A mutants predicted to be impaired in filament disassembly and then quantitatively evaluated the distribution and turnover of endogenous NM2B.

### NM2 dynamics is primarily regulated by its C-terminal tail

We evaluated the relative contribution of the motor and the rest of the heavy chain to the ability of NM2A to redistribute endogenous NM2B in COS7 cells using the NM2BA chimera (Baird et al., 2017), which comprises the NM2B motor domain fused to the NM2A rod, and comparing its effects to those produced by wild type NM2A or GFP alone as control (Figure 1).

**Figure 1.**
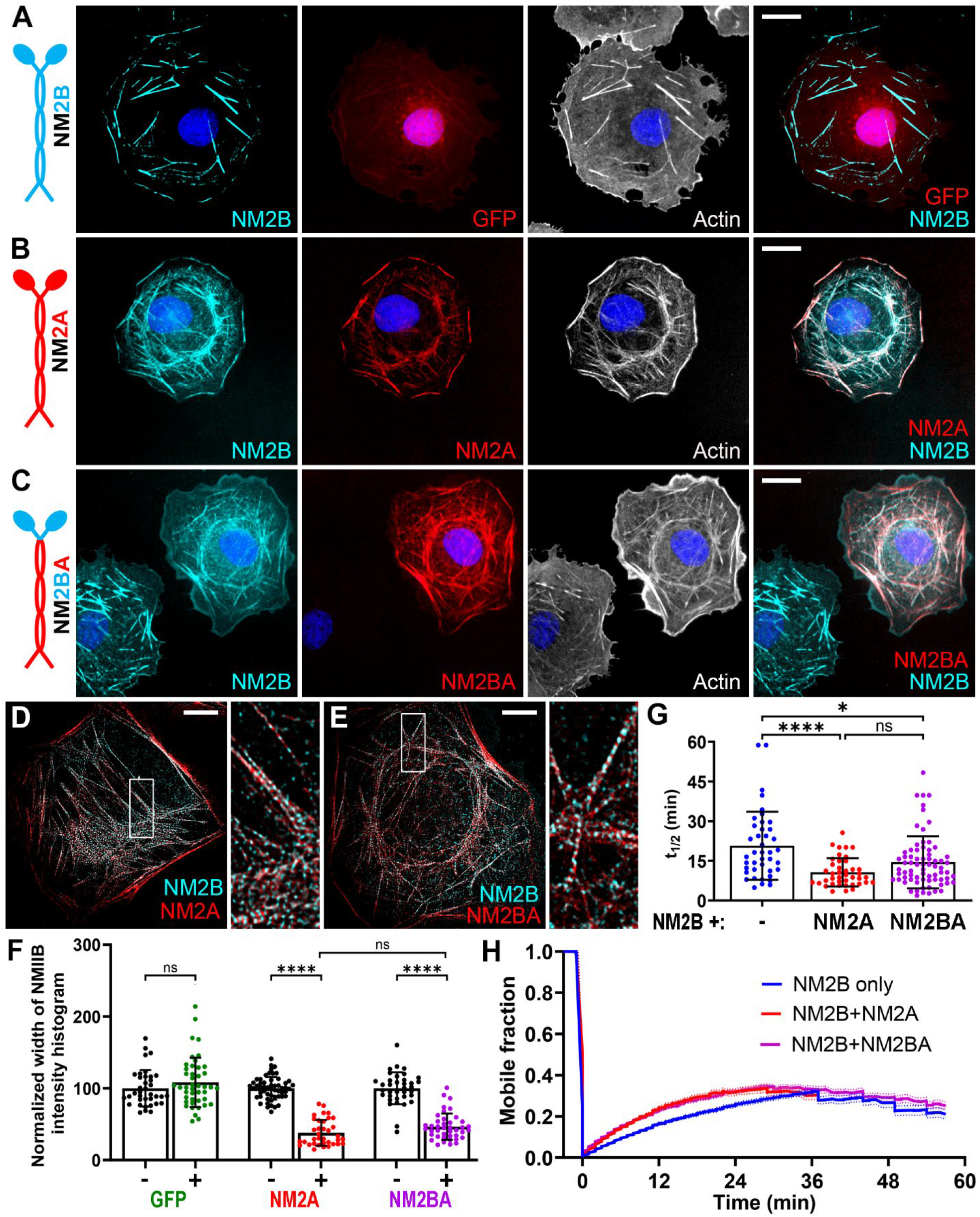
NM2 constructs with the NM2A tail reorganize and dynamize wild type NM2B. **(A-E)** Confocal (A-C) and DeepSIM (D, E) images of COS7 cells expressing indicated GFP-tagged constructs (red) and stained with NM2B antibody (cyan), phalloidin (A-C, white) and DAPI (A-C, blue). Diagrams of NM2 constructs are shown at left. Boxed regions in D and E are zoomed at right. Scale bars, 20 μm (A-C) and 10 μm (D, E). **(F)** Widths of NM2B immunofluorescence intensity histograms in cells expressing indicated constructs (+) after normalization against non-transfected cells (-) in the same samples. Dots, individual cells; bars, means; error bars, s.d.; ****, p<0.0001; ns, non-significant. **(G, H)** Results of FRAP assays showing halftimes of recovery (G) and average FRAP curves (H) of mCherry-NM2B in cells with no coexpression (blue), or with coexpression of GFP-NM2A (red) or GFP-NM2BA (magenta). Red curve is shorter because of significant remodeling of stress fibers after ∼30 min in the presence of NM2A. Error bars, s.d. (G) or s.e.m. (H); bars (G), mean; ****, p<0.0001; *, p<0.05; ns, non-significant.

Qualitatively, GFP-expressing cells retained the typical pattern of isolated NM2B-enriched ventral stress fibers, whereas overexpression of either GFP-NM2A or GFP-NM2BA reorganized this system into interconnected stress fibers with more homogeneous distribution of endogenous NM2B (Figure 1A-C). Using superresolution structured illumination microscopy (SIM), we observed efficient copolymerization of both NM2A and NM2BA with endogenous NM2B (Figure 1D, E). These copolymers appear as alternating dots corresponding to immunolabeled C-terminal tails of NM2B (color-coded cyan), which localize at the bare zone of bipolar filaments, and to GFP-tagged motor domains of ectopic constructs (color-coded red) positioned at bipolar filament ends.

We quantified the NM2B distribution by measuring the widths of NM2B immunofluorescence intensity histograms (Shutova et al., 2017), which become narrower for more homogeneous distributions of NM2B. Since the degree of NM2B redistribution depends on the expression level of the exogenous construct (Shutova et al., 2017) (Supplemental figure S1A), we assessed only highly expressing cells. The average histogram width in expressing cells normalized against the average value in non-expressing cells from the same coverslip was not changed by the expression of GFP but was significantly reduced by either GFP-NM2A or GFP-NM2BA expression (Figure 1F). These results suggest that the C-terminal tail of NM2A is primarily responsible for redistribution of NM2B. However, a non-significant trend toward greater values in the case of NM2BA relative to NM2A might reflect a marginal effect of the NM2A motor domain, the idea supported by experiments below.

As a complementary test, we used fluorescence recovery after photobleaching (FRAP) to determine whether expression of GFP-NM2BA, similar to GFP-NM2A (Shutova et al., 2017), accelerates the dynamics of wild type NM2B. We coexpressed mCherry-NM2B, as a reporter construct, and GFP-NM2B, GFP-NM2A or GFP-NM2BA, as test constructs (Figure 1G, H). The recovery curves of mCherry-NM2B were obtained from kymographs generated along the line crossing the bleached stress fiber and fitted to single exponential (Supplemental Figure S2A-C). The average halftime of recovery (t1/2) of mCherry-NM2B in stress fibers not undergoing significant remodeling (contracting, fusing, splitting, breaking, etc.) during 30-60 min imaging was 20.7 ± 12.9 min (mean ± s.d.) in the presence of GFP-NM2B as control, but was significantly reduced by either GFP-NM2A (10.7 ± 5.4 min) or GFP-NM2BA (14.5 ± 9.8 min). Although not significantly, the effect of NM2BA appeared weaker than that of NM2A, hinting again that the NM2A motor may contribute to the phenotype. On the other hand, the NM2B mobile fractions did not show significant differences among these conditions (Supplemental figure S1B).

Together, these results show that the ectopically expressed NM2BA chimera, similar to wild type NM2A, can redistribute and dynamize endogenous NM2B in COS7 cells, thus highlighting the key role of the NM2A tail in these effects. These findings are consistent with the existing literature indicating that NM2A’s own dynamics is largely controlled by its C-terminal tail (Sandquist and Means, 2008; Taneja and Burnette, 2019; Vicente-Manzanares et al., 2008).

### NHT and C-terminal phosphorylation of NM2A cooperate in regulating NM2B in trans

The NHT and the C-terminal phosphorylation sites play key roles for NM2A turnover; when these disassembly mechanisms were disabled, the dynamics of such NM2A mutants in vitro and in cells was reduced (Breckenridge et al., 2009; Dulyaninova et al., 2007; Dulyaninova et al., 2005; Rai et al., 2017) and their coassembly with NM2B in cells was increased (Weissenbruch et al., 2022).

We tested the roles of these disassembly mechanisms for redistribution of endogenous NM2B using NM2A mutants with impaired dynamics; namely, NM2A with truncated NHT (NM2A-ΔNHT), NM2A with three phosphorylation-blocking substitutions (S1943A in the NHT and S1916A and S1915A just upstream of the NHT; NM2A-3SA), as well as a combination of these mutations (NM2A-ΔNHT/2SA) (Figure 2). The phosphorylation of S1943 also prevents association of S100A4, an NM2A disassembly factor (Dulyaninova et al., 2005; Ford et al., 1997; Li et al., 2003). Thus, the NM2A-3SA mutation is expected to disable both phosphorylation-dependent and S100A4-dependent mechanisms of NM2A disassembly. Of note, S100A4 is not expressed in COS7 cells (Li et al., 2010), but other S100 family members (Ecsedi et al., 2018) may act in such capacity in COS7 cells.

**Figure 2.**
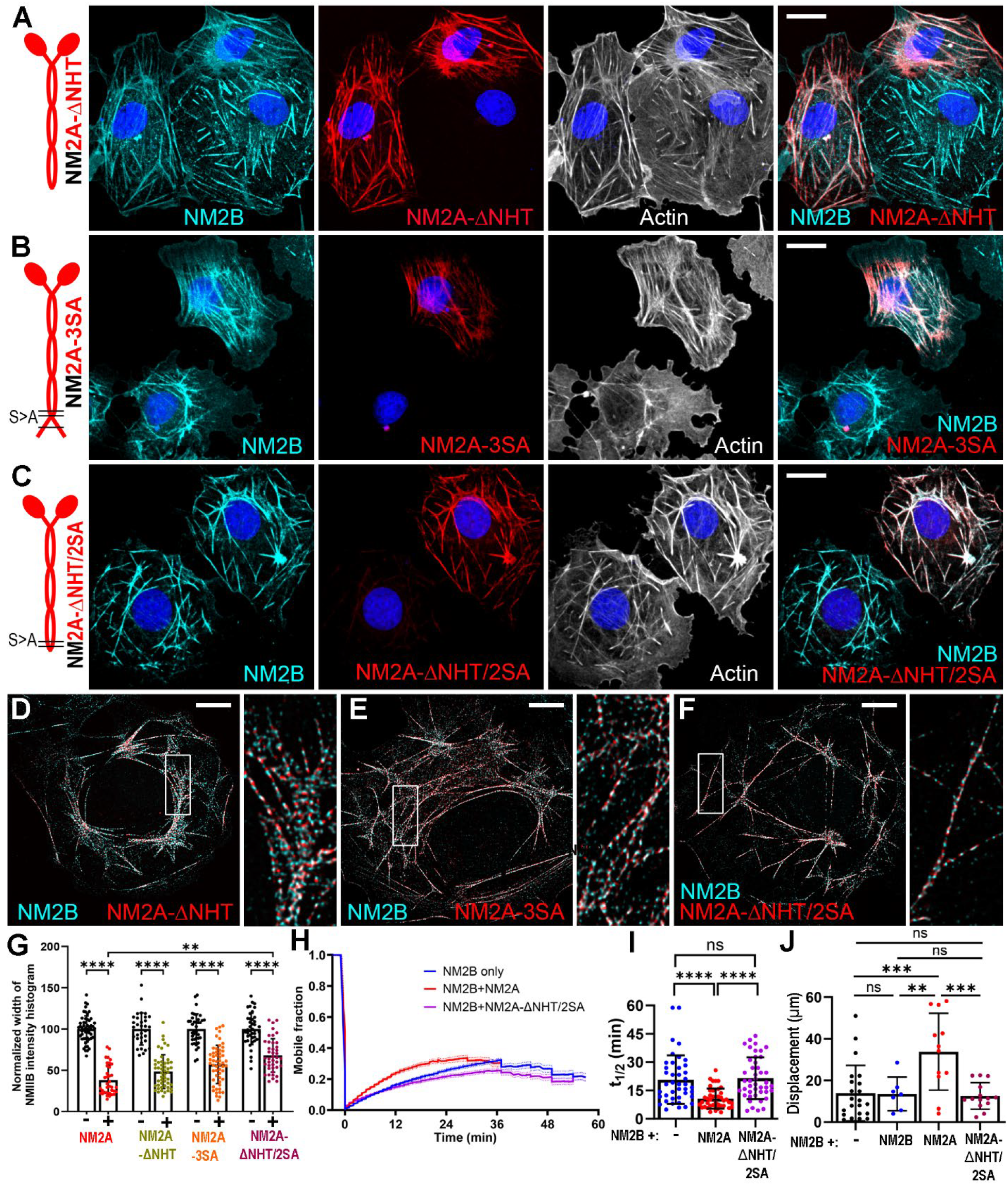
NHT and C-terminal phosphorylation sites of NM2A jointly regulate NM2A-dependent NM2B dynamics and cell motility. **(A-F)** Confocal (A-C) and SIM (D-F) images of COS7 cells expressing indicated GFP-tagged constructs (red) and stained with NM2B antibody (cyan), phalloidin (A-C, white) and DAPI (A-C, blue). Diagrams of NM2 constructs are shown at left. Boxed regions in D-F are zoomed at right. Scale bars, 20 μm (A-C) and 10 μm (D-F). **(G)** Widths of NM2B immunofluorescence intensity histograms in cells expressing indicated constructs (+) after normalization against non-transfected cells (-) in the same samples. Data for ± NM2A are repeated from figure 1F for comparison. Dots, individual cells; bars, means; error bars, s.d.; ***, p<0.0001; **, p<0.01. **(H, I)** Results of FRAP assays showing average FRAP curves (H) and halftimes of recovery (I) of mCherry-NM2B in cells with no coexpression (blue), or with coexpression of GFP-NM2A (red) or GFP-NM2BA (magenta). Data for NM2B only and NM2B+NM2A are repeated from figure 1G and 1H for comparison. Error bars, s.e.m. (H) and s.d. (I); bars (I), mean; ****, p<0.0001; ns, non-significant. **(J)** Average net displacement of COS7 cells expressing the indicated constructs over a 14-h time period. Dots, individual cells; bars, means; error bars, s.d.; ***, p < 0.001, **, p<0.01; ns, not significant.

When expressed in COS7 cells, both NM2A-ΔNHT and NM2A-3SA efficiently copolymerized with the endogenous NM2B, as seen by both confocal microscopy (Figure 2A, B) and SIM (Figure 2D, E). Surprisingly, these mutants still efficiently redistributed NM2B (Figure 2A, B) and significantly reduced the NM2B histogram widths, as compared with non-expressing cells (Figure 2G). Yet, given a trend toward wider histograms induced by each mutant than by wild type NM2A, these mutations may have a slight inhibitory effect on driving the NM2B redistribution. The combination of both mutations (NM2A-ΔNHT/2SA) had a significantly stronger negative effect on NM2A-driven NM2B redistribution than individual ΔNHT or 3SA mutations, as the average histogram width of NM2B in the presence of NM2A-ΔNHT/2SA was significantly greater than that in the presence of NM2A, but still significantly lower than in control untransfected cells, showing that the redistribution was still incomplete (Figure 2G). These results suggest that the NHT and the three C-terminal phosphorylation sites cooperatively control the NM2A turnover, which enables NM2A to redistribute NM2B in trans, but that their collective incapacitation is not sufficient to abrogate the NM2A-dependent NM2B redistribution.

By FRAP (Figure 2H, I; Supplemental Figure 2D), the halftime of NM2B recovery was not significantly reduced in the presence of NM2A-ΔNHT/2SA (21.5 ± 11.0 min). Thus, the two tests showed apparently conflicting results: NM2A-ΔNHT/2SA still can redistribute endogenous NM2B albeit less efficiently than NM2A (the histogram test) but, in contrast to NM2A, could not dynamize the NM2B turnover (the FRAP test). A possible explanation is the different timing of the two assays. Perhaps, the ΔNHT/2SA mutation can sufficiently impair the NM2A filament disassembly to make the phenotype detectable over long term in the redistribution test (2 days), when extensive restructuring of the contractile system likely takes place, but not in the short term FRAP assay (30-60 min) when NM2 molecules come in and out of a static stress fiber.Given the minimal effects produced by NM2A-ΔNHT and NM2A-3SA in the redistribution test, we did not assay these mutants by FRAP.

We next analyzed the impact of the ΔNHT/2SA mutation in NM2A, and therefore of NM2A turnover, on COS7 motility using live cell imaging. Expression of NM2A, as expected (Shutova et al., 2017), significantly increased COS7 cell migration, as compared to wild-type COS7 cells, while ectopic expression of NM2B had no effect. Importantly, the overexpressed NM2A-ΔNHT/2SA mutant failed to enhance cell motility (Figure 2J). These findings show that the ability of NM2A to enhance cell migration depends on its own turnover, which in turn requires synergistic inputs from NM2A’s NHT and S1915/S1916 phosphorylation sites.

### The NM2A motor domain plays an important role in accelerating NM2B dynamics in trans

A significant, but incomplete, inhibition of NM2A’s ability to redistribute endogenous NM2B by the ΔNHT/2SA mutation suggests that additional features of the NM2A molecule, either elsewhere in the NM2A tail or in its motor domain, also contribute. We tested the role of the NM2A motor domain by introducing the ΔNHT, 3SA, or ΔNHT/2SA mutation into the NM2BA chimera (Figure 3). All these constructs successfully copolymerized with endogenous NM2B (Figure 3A-F), although SIM showed that NM2BA-ΔNHT/2SA did so less efficiently (Figure 3F). However, in the redistribution test (Figure 3A-C, G), only NM2BA-ΔNHT slightly reduced the histogram width with minimal significance, whereas NM2BA-3SA and NM2BA-ΔNHT/2SA did not cause significant changes. However, the NM2BA-ΔNHT/2SA mutant showed a trend of increasing rather than decreasing the histogram width, suggesting that it could be even less dynamic than wild type NM2B.

**Figure 3.**
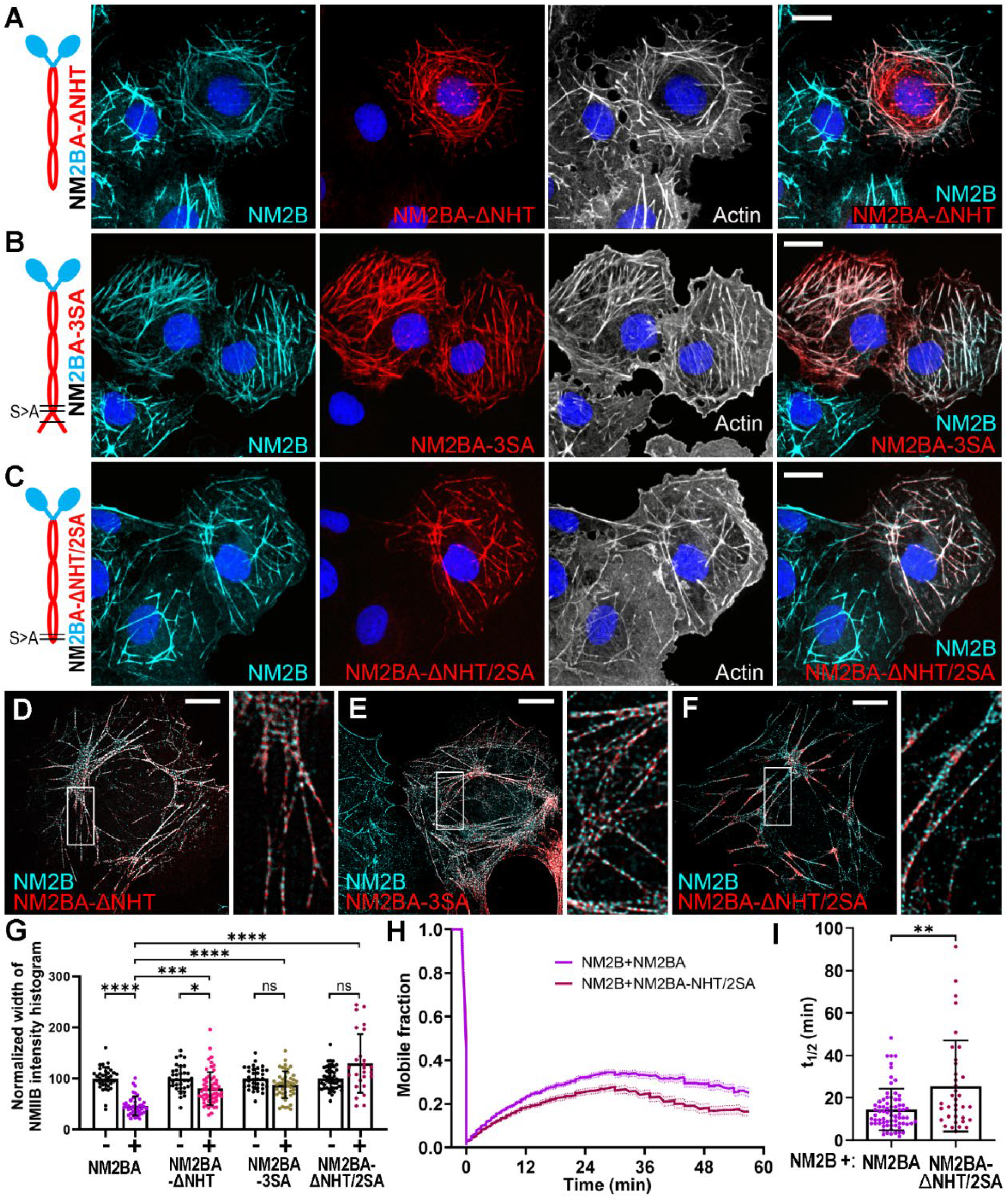
NM2A motor domain contributes to acceleration of NM2B dynamics in trans **(A-F)** Confocal (A-C) and DeepSIM (D-F) images of COS7 cells expressing indicated GFP-tagged constructs (red) and stained with NMIIB antibody (cyan), phalloidin (A-C, white) and DAPI (A-C, blue). Diagrams of NM2 constructs are shown at left. Boxed regions in D-F are zoomed at right. Scale bars, 20 μm (A-C) and 10 μm (D-F). **(G)** Widths of NM2B immunofluorescence intensity histograms in cells expressing indicated constructs (+) after normalization against non-transfected cells (-) in the same samples. Data for ±NM2BA are repeated from figure 1F for comparison. Dots, individual cells; bars, means; error bars, s.d.; ****, p<0.0001; ***, p<0.001; *, p<0.05; ns, non-significant. **(H, I)** Results of FRAP assays showing average FRAP curves (H) and halftimes of recovery (I) of mCherry-NM2B in cells coexpressing GFP-NM2A (magenta) or GFP-NM2BA (dark red). Data for NM2B+NM2BA are repeated from figure 1G and 1H for comparison. Error bars, s.e.m. (H) or s.d. (I); bars (I), mean; **, p<0.01.

In the FRAP assay (Figure 3H, I; Supplemental Figure 2E), the NM2BA-ΔNHT/2SA expression significantly increased t1/2 of NM2B recovery (25.5 ± 21.5 min) relative to NM2BA (14.5 ± 9.8 min), but it was comparable to t1/2 of NM2A-ΔNHT/2SA. Therefore, when the turnover-impaired NM2A tail is linked to the NM2B motor, such chimera is not able to either redistribute or dynamize NM2B in trans.

Thus, the presence of NM2A motor allows the turnover-impaired NM2A-ΔNHT/2SA mutant to redistribute NM2B during long-term cytoskeletal reorganization but not to accelerate the short-term exchange of NM2B molecules in the stress fiber. One explanation is that once released from a stress fiber (by NM2 tail-dependent mechanisms or by actin disassembly if the latter is impaired), NM2 monomers or filaments, including heteropolymers, can efficiently move along actin filaments to distant places if they possess the NM2A motor but less efficiently if they have only NM2B motors. Unfolded, and therefore motor-active, NM2 monomers are indeed present in cells (Shutova et al., 2014), while processive movement of NM2 filaments was also reported (Melli et al., 2018; Vitriol et al., 2023).

### Mechanism(s) regulating NM2B dynamics

If NM2A redistributes NM2B due to its own fast turnover, then accelerating dynamics of ectopic NM2B should have a similar effect on endogenous wild type NM2B. Since S1935 makes the highest contribution to the regulation of NM2B dynamics (Juanes-Garcia et al., 2015), we introduced the phosphomimetic S1935D (SD) mutation into the NM2B tail of NM2B or NM2AB to accelerate their own turnover and potentially allow them to redistribute endogenous wild type NM2B similar to the NM2 constructs containing the NM2A tail.

For these experiments, however, we could not use immunostaining of endogenous NM2B, because NM2B antibodies recognize the C-terminus of the NM2B heavy chain, which is also a part of the expressed constructs in this set of experiments. Therefore, we tagged the endogenous NM2B heavy chain with mScarlet (NM2B-mScarlet) by CRISPR/Cas9 technology and used this knockin COS7 cell line to overexpress GFP-tagged NM2B, NM2B-SD, NM2AB or NM2AB-SD (Figure 4). These constructs successfully copolymerized with endogenous NM2B-mCherry (Figure 4A-H).

**Figure 4.**
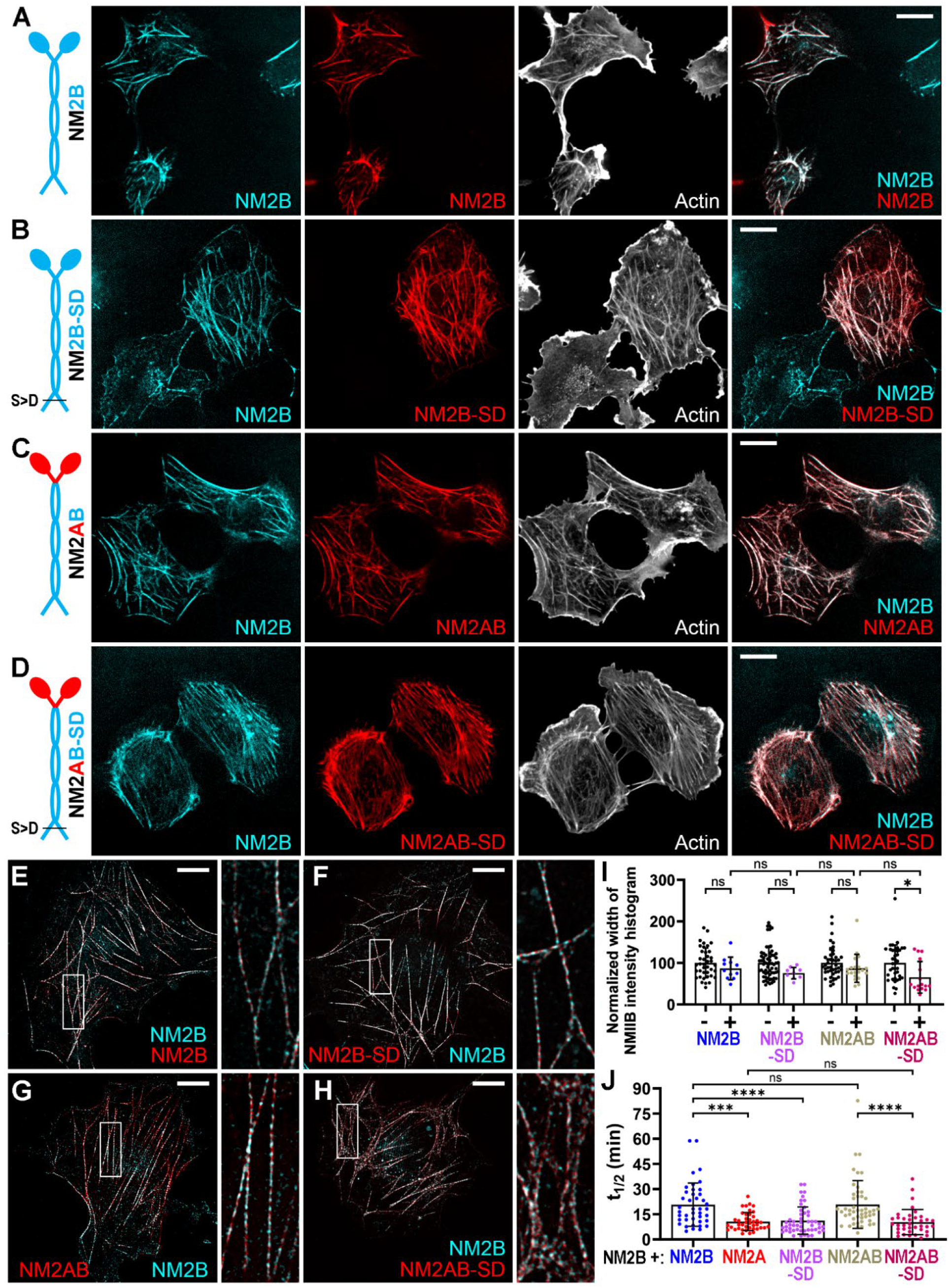
Effects of dynamizing mutation in the NMIIB tail depend on the motor **(A-H)** Confocal (A-D) and DeepSIM (E-H) images of COS7 cells expressing indicated GFP-tagged constructs (red) and endogenously tagged NM2B-mScarlet (cyan) and phalloidin (white). Boxed regions in E-H are zoomed at right. Scale bars, 20 μm (A-D) and 10 μm (E-H). **(I)** Widths of NM2B immunofluorescence intensity histograms in cells expressing indicated constructs (+) after normalization against non-transfected cells (-) in the same samples. Dots, individual cells; bars, means; error bars, s.d.; *, p<0.05; ns, non-significant. **J)** Halftimes of mCherry-NM2B recovery in the presence of indicated constructs. ***, p<0.001; ****, p<0.0001; ns, non-significant.

Expression of wild type GFP-NM2B did not redistribute NM2B-mScarlet in knockin COS7 cells (Figure 4A, I), as expected (Shutova et al., 2017). Surprisingly, GFP-NM2B-SD also did not have a significant effect by this test (Figure 4B, I), but was able to dynamize NM2B-mScarlet by the FRAP assay (Figure 4J, Supplemental Figure 3A, D). Thus, the S1935D mutation, as predicted, dynamizes endogenous NM2B-mScarlet by promoting disassembly of the heterotypic GFP-NM2B-SD/NM2B-mScarlet filaments. However, due to the slow NM2B motor, dissociated monomers might not be able to significantly move away from the site of disassembly preventing large scale redistribution. Consistent with this idea, the impaired NM2B motor activity was shown to further reduce the already slow intrinsic turnover of NM2B (Balaban et al., 2023; Wang et al., 2023).

The results were different in the case of the NM2AB chimera. With an intact tail, GFP-NM2AB neither redistributed endogenous NM2B-mScarlet (Figure 4C, I) nor affected its recovery by FRAP (Figure 4J, Supplemental Figure 3B, E). However, the NM2AB-SD chimera was efficient in both tests: it reduced both an average histogram width of NM2B-mCherry distribution indicating significant redistribution (Figure 4D, I) and the t1/2 of NM2B-mScarlet recovery (Figure 4J, Supplemental Figure 3C, E), thus showing significant dynamizing activity. These data suggest that although S1935 phosphorylation plays an important role in the local NM2B turnover, it is not sufficient to cause large-scale NM2B redistribution unless linked to fast NM2A motor. On the other hand, the presence of the NM2A motor, if it is linked to the intrinsically slowly turning over NM2B tail, is insufficient to redistribute endogenous NM2B in trans.

## Conclusion

Previous studies addressing the regulation of NM2 dynamics revealed key roles of the NHT, phosphorylation and protein-protein interactions targeting the C-terminal region of NM2 heavy chains (Chinthalapudi and Heissler, 2024; Dulyaninova and Bresnick, 2013; Shutova and Svitkina, 2018). However, the discovery of copolymerization of different NM2 paralogs added more complexity to this question (Beach et al., 2014; Shutova et al., 2014). We previously formulated a model of self-organization of the contractile system in cells, wherein cycles of nonselective copolymerization of NM2A and NM2B paralogs combined with preferential dissociation of NM2A subunits leads to different outcomes that depend on NM2A/NM2B ratios in cells (Shutova et al., 2017; Shutova and Svitkina, 2018). A key aspect of this model is the reciprocal influence of different NMII paralogs on each other’s dynamics.

In this study, we demonstrated that the known mechanisms regulating NM2A’s own dynamics are also responsible for its ability to influence the dynamics of NM2B in cells, thus providing an experimental validation of our model. Furthermore, we revealed the specific features of the NM2A heavy chain that enable such acceleration of NM2B dynamics in trans. They include the NHT, phosphorylation of S1915, S1916, and/or S1943 residues, and surprisingly, the motor domain. Moreover, these individual mechanisms act synergistically. Even though each of them contributes to fast NM2A dynamics, collectively, they have a greater effect. These effects are not limited to NM2A-mediated regulation of NM2B dynamics but are also responsible for remodeling the actin cytoskeleton during cell movement and for the efficiency of cell migration. The observed effect on cell migration is consistent with our model of NM2A/2B self-sorting, in which at the optimal NM2A/2B, these paralogs form a polarized gradient essential for front-rear polarization and directed cell migration (Shutova et al., 2017; Shutova and Svitkina, 2018). In this mechanism, NM2A uses both its fast motor and rapid filament turnover to efficiently redistribute throughout the cell, including the vicinity of the leading edge, to support cell adhesion. At the same time, NM2B due to its longer duty ratio, slower motor-driven movement, and less efficient filament turnover, drift away from the edge with the actin retrograde flow and accumulates in the cell posterior to retract the cell rear and prevent unproductive protrusions there.

## MATERIALS AND METHODS

### Cell Culture

COS7 green monkey kidney cells were cultured in high-glucose GlutaMAX-containing DMEM (Gibco) supplemented with 10% fetal bovine serum (FBS, Gibco), and 1% penicillin/streptomycin at 37°C in a humidified incubator with 5% CO_2_. For transfection, cells were seeded onto 35-mm culture dishes and allowed to adhere overnight before transfection. Cells were periodically tested for mycoplasma contamination (MycoStrip, InvivoGen).

The COS7 NMIIB:3xGGGGS-mScarlet knockin cell line was generated via CRISPR/Cas9-mediated in-frame insertion of a 3xGGGGS-mScarlet linker into exon 41 of the MYH10 locus.

The knockin was performed in wild-type COS-7 cells by Ubigene Biosciences Co., Ltd. (Guangzhou, China).

### Plasmids and Transfection

The following plasmids were obtained from Addgene: CMV-GFP-NMHC II-A (#11347), CMV-GFP-NMHC II-B (#11348), mCherry-MyosinIIB-C-18 (#55106), pCMV-eGFP-NMHC-IIAΔtailpiece (#35689), pEGFP-NMHC-IIA-3xA (#101041), and mEGFP-C1 (#54759). Chimeric constructs GFP-NM2A2B and GFP-NM2B2A were kindly provided by Dr. Clare Waterman (Baird et al., 2017).

Deletions and point mutations were introduced using the Q5 Site-Directed Mutagenesis Kit (New England Biolabs). The list of PCR primers (IDT) is provided in Table S1. The NM2A-ΔNHT/2SA mutant, which lacks the NHT (amino acids 1928–1961) and carries S1915A and S1916A substitutions, was generated from pCMV-eGFP-NMHC-IIAΔtailpiece (#35689).

Mutants of chimeric constructs, NM2BA-ΔNHT, NM2BA-3SA (S1915A, S1916A, S1943A), and NM2BA-ΔNHT/2SA were generated from the GFP-NM2B2a plasmid. The S1935D point mutation was introduced into CMV-GFP-NMHC II-B and GFP-NM2A2B yielding NM2B-S1935D and NM2AB-S1935D, respectively. All constructs were validated by in-house sequencing at the Penn Genomics and Sequencing Core, and primers were synthesized by Integrated DNA Technologies (IDT).

Transfections were performed using Lipofectamine 3000 (Invitrogen) following the manufacturer’s instructions.

### Immunofluorescence staining

Twenty-four hours post-transfection, cells were replated onto glass coverslips for 24 hours and fixed by 4% formaldehyde in PBS for 20 min at room temperature and subsequently permeabilized with 0.03% Triton X-100 in PBS. Immunostaining for NM2B was performed as described (Shutova et al., 2017) using a rabbit polyclonal NM2B antibody (#3404; Cell Signaling) and Alexa Fluor-568 anti-rabbit IgG secondary antibody (#A-11011, Invitrogen). F-actin was labeled with Alexa Fluor-647 phalloidin (#8940; Cell Signaling) or phalloidin-iFluor 405 (#176752, Abcam). Coverslips were mounted with ProLong Gold antifade reagent (#36935, Invitrogen) containing DAPI for nuclear staining.

#### Confocal Microscopy

Z-stack images were acquired using Eclipse Ti-U inverted microscope (Nikon Instruments, Japan) equipped with LUNV 7-line laser launch (Nikon), CSU-X1 spinning disk (Yokogawa), QuantEM:512SC CCD camera (Photometrics), and Plan Apo 60×1.4 NA oil-immersion objective and operated by Nikon Imaging Software (NIS-Elements). Excitation was performed using 405-nm, 488-nm, 561-nm, and 640-nm laser lines along with a quad-bandpass filter for simultaneous detection of DAPI, GFP, Alexa Fluor-568 (anti-NM2B), and Alexa Fluor-647 phalloidin fluorescence, respectively.

### Fluorescence recovery after photobleaching (FRAP)

For FRAP experiments, COS7 cells were cotransfected with various GFP-tagged NM2 constructs along with mCherry-NM2B. Twenty-four hours post-transfection, cells were replated onto glass-bottom 35-mm Petri dishes and allowed to spread for 6 hours. Prior to imaging, cells were transferred to CO_2_-independent L15 medium for 1 hour at 37°C. During live cell imaging cells were maintained at 37°C in stage-top incubation chamber (Okolab USA Inc.). Single confocal images of GFP and mCherry channels were acquired to record transfections. Then, after time-lapse acquisition of 10 pre-bleach frames of mCherry-NM2B, rectangular regions (∼ 16x13 µm) were bleached for 5 seconds using a 405-nm laser at 100% power. Subsequent imaging of NM2B-mCherry was conducted at a single confocal plane using 60×1.4 NA lens at 28.7-second intervals per frame for 1 hour.

### Cell motility

Live-cell imaging of COS7 cells expressing with various ectopic NM2 constructs was performed 24 hours post-transfection. Cells were seeded onto four-well chamber coverglass (#155382, Nunc Lab-Tek II), allowed to spread for 6 hours at 37°C and were incubated with Hoechst dye (#33342, Thermo Scietific) according to the manufacturer’s instructions to label nuclei prior to confocal time-lapse imaging, which was carried out using a Plan Apo 20× 0.75 NA objective and Eclipse Ti-U inverted microscope described above. Images were captured every 10 minutes for 14 hours using 405-nm and 488-nm laser lines and a quad-bandpass emission filter to simultaneously detect Hoechst and GFP signals.

### Structured Illumination Microscopy (SIM)

Structured illumination microscopy (SIM) of fixed cells was performed using CrestOptics X-Light DeepSIM standalone module mounted on a Nikon Eclipse Ti2E inverted microscope (Nikon Instruments, Japan) equipped with sCMOS camera (Kinetix), CSU-X1 spinning disk (Yokogawa), and Plan Apochromat λD 100×NA 1.45 oil-immersion objective, providing a lateral pixel size of 100 nm (XY). Imaging was carried out using a 488-nm excitation laser with a 510/40 emission filter for GFP and a 545-nm excitation laser with a 595/50 emission filter for NM2B-mScarlet and Alexa Fluor-568. After acquisition of Z-stack images, image reconstruction was performed using Nikon Imaging Software (NIS-Elements).

### Image analysis

#### Distribution of endogenous NM2B

The distribution of immunolabeled endogenous NM2B in COS7 cells was quantified using ImageJ by measuring the widths of the NM2B immunofluorescence intensity (in wild type COS7 cells) or mScarlet fluorescence intensity (in COS7 knockin cells) histogram (1-99%) in 16-bit images, as described previously (Shutova et al., 2017). Only transfected cells with total fluorescence intensity above a threshold roughly equivalent to an average median value among different conditions were included in the analysis to ensure sufficient phenotypic effect upon expression of tested constructs. The values obtained from transfected cells were normalized against the average width of NM2B histogram in non-transfected cells in the same samples and expressed as percentages. Data were collected from three independent experiments, each including both transfected and non-transfected cells.

#### FRAP analysis

A kymograph along a line (10-30 pixel wide, equivalent to 2.2-6.6 μm) crossing the bleached stress fiber was generated using the KymoResliceWide plugin in ImageJ. The NM2B intensity profiles from the kymographs were obtained along a 3-pixel wide (0.67 μm) segmented line using Plot Profile function in ImageJ. Analysis of these profiles was performed using the Stowers plugin suite in ImageJ. First, the profiles were normalized to the 0 - 1 range using “normalize trajectories jru v1” plugin and fitted to a single exponential using the “batch FRAP fit jru v1” plugin. Only the profiles that fit the single exponential within the 30-60 min time frame were used. The fit output included pre-FRAP, baseline, amplitude, and tau values. The half-time recovery (t_1/2_) in minutes was calculated by converting frames to minutes and multiplying tau by 0.69. The mobile fraction was determined using the formula: amplitude/(prefrap-baseline).

#### Cell migration analysis

Time-lapse sequences of the nuclear motility (Hoechst channel) were processed using ImageJ by applying bleach correction (histogram matching), background subtraction (50-pixel rolling ball radius), Gaussian blur (SigmaRadius = 3), and adjusting brightness/contrast. The processed images were converted to 8-bit and analyzed using the TrackMate plugin in ImageJ. Tracks were generated using the DoG detector (25 µm diameter) and Simple LAP tracker (linking and gap-closing distance: 50 µm; max gap: 2 frames). Tracks were manually verified using a composite image overlay to confirm GFP expression. Track displacement in microns was extracted from validated tracks and compared across conditions. Non-expressing wild-type COS7 cells served as control.

#### Statistics

Statistical comparisons were performed using the Kruskal-Wallis test in GraphPad Prism 10.2.2. The same software was used to generate graphs.

## ONLINE SUPPLEMENTAL MATERIAL

Figure S1 shows correlations between histogram widths of endogenous NM2B and the expression levels of ectopic constructs and FRAP mobile fractions of NM2B in different conditions. Figures S2 and S3 show examples of FRAP of NM2B in stress fibers in cells expressing constructs containing NM2A or NM2B tails, respectively. Table S1 gives a list of primers used for mutagenesis.

## DATA AVAILABILITY

All other data generated or analyzed for this study are presented in the manuscript and its supplementary information files.

## ACKNOWLEDGMENTS

We thank Dr. Clare Waterman (NIH) for providing NM2 chimeras, Justin Bi and Aditya Saha for help with data analyses, and Changsong Yang and Xingyuan Fang for useful discussions. This work is supported by NIH grant R35 GM 140832 and NSF grant CMMI-1548571 to TS.

## Notes

### Competing Interest Statement

The authors have declared no competing interest.

