## Supplementary material for "Nonmuscle myosin 2 turnover in cells is synergistically controlled by the tail and the motor domain"

SUPPLEMENTAL MATERIALS

**Table S1:** PCR primers used for mutagenesis.

| Constructs | Primer sequence |
|---|---|
| NM2A-ΔNHT/2SA | Fw 5'-atgaaccgcgaagtcgccgccctaaagaacaagc-3' |
| NM2BA-ΔNHT/2SA | Rv 5'-ggcatcggccgtctcagtg-3' |
| NM2BA-S1915/16A | Fw 5'-atgaaccgcgaagtcgccgccctaaagaacaagc-3' |
| | Rv 5'-ggcatcggccgtctcagtg-3' |
| NM2BA-S1943A | Fw 5'-gccggggatggcgccgacgaagagg-3' |
| | Rv 5'-gcctttccgggccattcggc-3' |
| NM2BA-ΔNHT | Fw 5'-ccgggatccaccggatctagataac-3' |
| | Rv 5'-gcccgcggtaccttacggcaggtccccgcg-3' |
| NM2B-S1935D | Fw 5'-adcsafsdfsdsdsdsed-3' |
| NM2AB-S1935D | Rv 5'-sfgfgfdgfdhfdhfg-3 |

SUPPLEMENTAL FIGURES

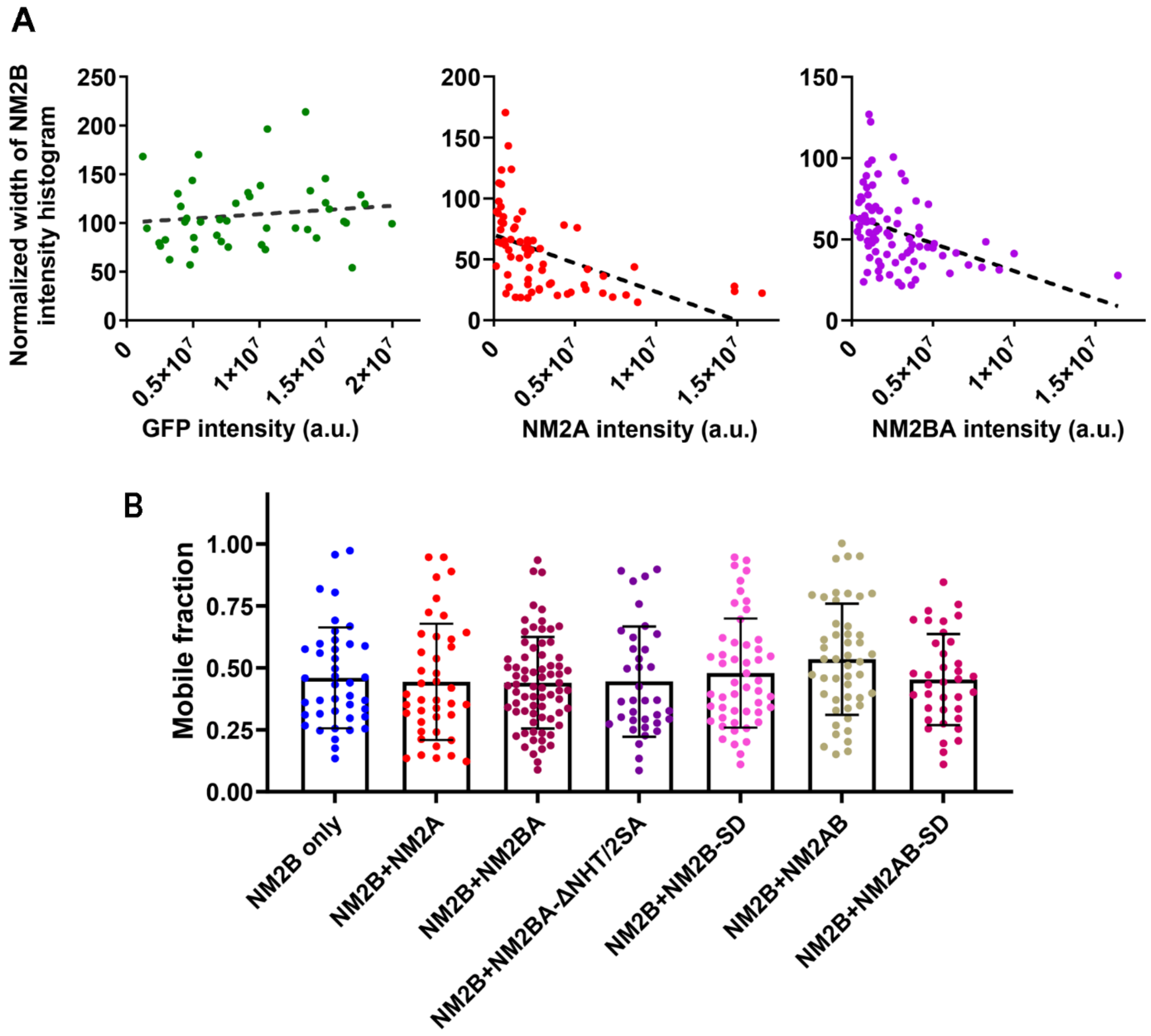


**Figure S1.** (A) Widths of NM2B immunofluorescence intensity histograms decrease with high levels of overexpression of GFP-NM2A or GFP-NM2BA, but of GFP alone. Dots, individual cells; dashed lines represent linear fits.

(B) Mobile fractions of mCherry-NM2B in stress fibers of cells expressing indicated constructs. Dots, individual cells; bars, means; error bars, s.d. No significant differences are found among these datasets.

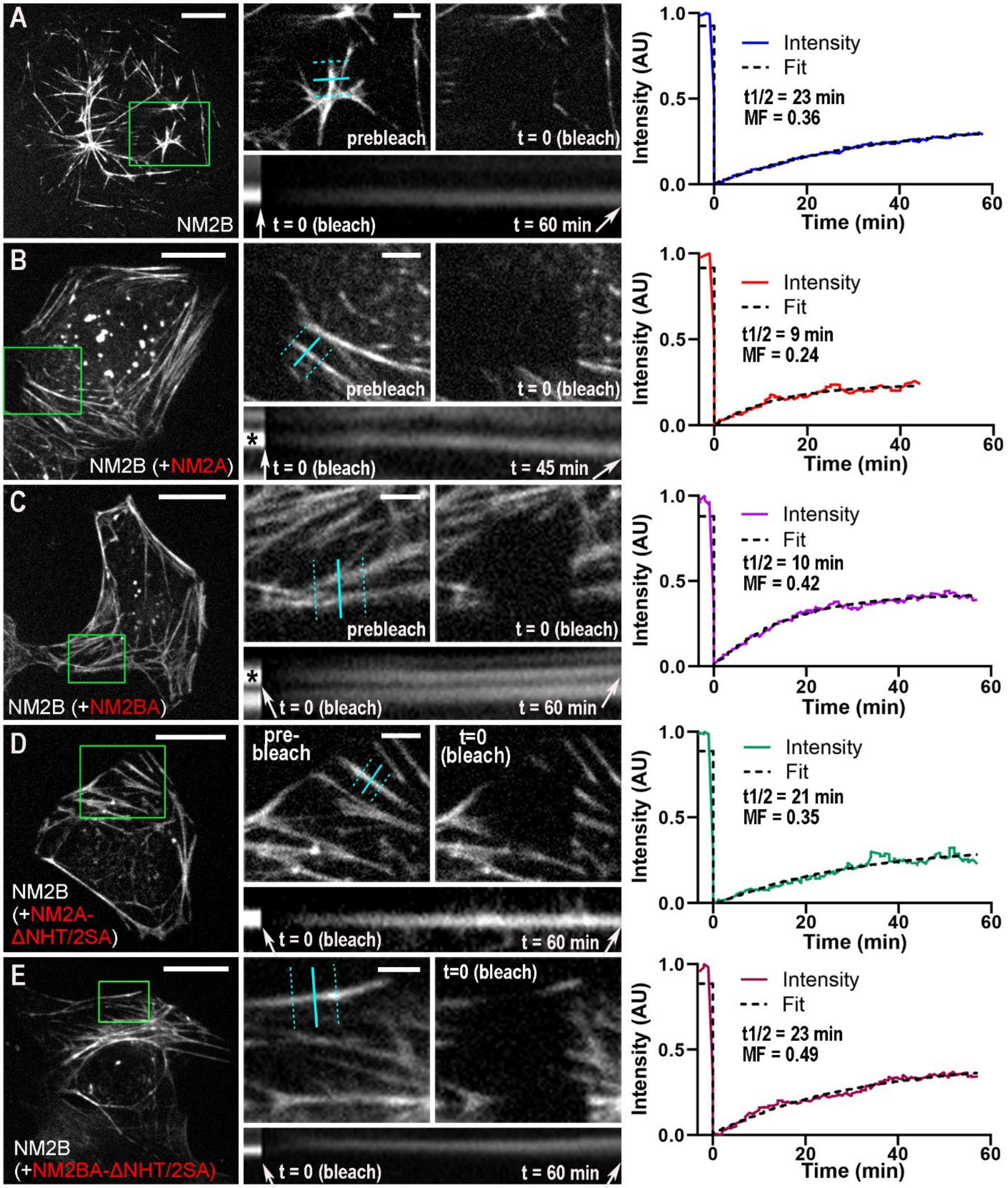


**Figure S2.** Examples of FRAP of mCherry-NM2B (white) in individual stress fibers in COS7 cells expressing mCherry-NM2B only (A), or together with GFP-NM2A (B), GFP-NM2BA (C), NM2A-ΔNHT/2SA (D), or NM2BA-ΔNHT/2SA (E). Green boxed regions in main panels are enlarged in the upper middle panels before and immediately after bleach. Kymographs (bottom middle) are taken along the cyan lines with intensities averaged along the segment between the dashed cyan lines. Graphs (right) show recovery curves (colored) and exponential fits (black dashed lines) for tested stress fibers (marked by asterisks in B and C). Halftimes (t1/2) and mobile fractions (MF) are shown for each plot. Scale bars, 20 μm (main panels) and 5 μm (enlargements).

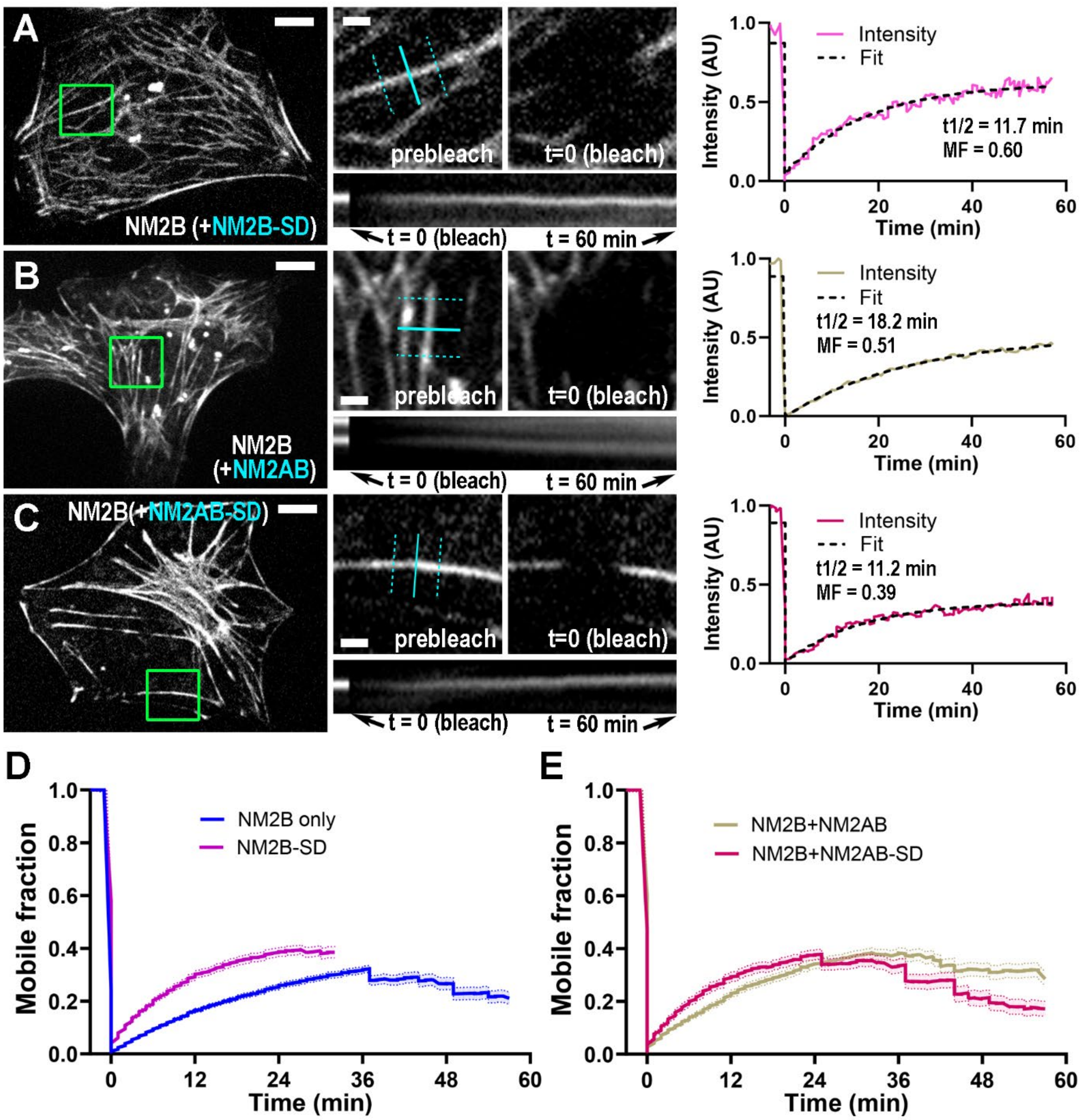


**Figure S3.** (A-C) Examples of FRAP of mCherry-NM2B (white) in individual stress fibers in COS7 cells coexpressing mCherry-NM2B with GFP-NM2B-SD (A), GFP-NM2AB (B), or NM2AB-SD (C). Green boxed regions in main panels are enlarged in the upper middle panels before and immediately after bleach. Kymographs (bottom middle) are taken along the cyan lines with intensities averaged along the segment between the dashed cyan lines. Graphs (right) show recovery curves (colored) and exponential fits (black dashed lines) for tested stress fibers. Halftimes (t1/2) and mobile fractions (MF) are shown for each plot. Scale bars, 20 µm (main panels) and 5 µm (enlargements).

(D, E) Average FRAP curves of mCherry-NM2B in cells coexpressing mCherry-NM2B with GFP-NM2B-SD (D), GFP-NM2AB (E) or GFP-NM2AB-SD (E). Error bars, s.e.m.
